# Detailed mechanisms for unintended large DNA deletions with CRISPR, base editors, and prime editors

**DOI:** 10.1101/2024.01.04.574288

**Authors:** Gue-ho Hwang, Seok-Hoon Lee, Minsik Oh, Segi Kim, Omer Habib, Hyeon-Ki Jang, Heon Seok Kim, Chan Hyuk Kim, Sun Kim, Sangsu Bae

## Abstract

CRISPR-Cas9 nucleases are versatile tools for genetic engineering cells and function by producing targeted double-strand breaks (DSBs) in the DNA sequence. However, the unintended production of large deletions (>100 bp) represents a challenge to the effective application of this genome-editing system. We optimized a long-range amplicon sequencing system and developed a k-mer sequence-alignment algorithm to simultaneously detect small DNA alteration events and large DNA deletions. With this workflow, we determined that CRISPR-Cas9 induced large deletions at varying frequencies in cancer cell lines, stem cells, and primary T cells. With CRISPR interference screening, we determined that end resection and the subsequent TMEJ [DNA polymerase theta-mediated end joining] repair process produce most large deletions. Furthermore, base editors and prime editors also generated large deletions despite employing mutated Cas9 “nickases” that produce single-strand breaks. Our findings reveal an important limitation of current genome-editing tools and identify strategies for mitigating unwanted large deletion events.

## Introduction

The CRISPR-Cas system, derived from a prokaryotic immune response, uses single-guide RNAs (sgRNAs) to recognize target DNA sequences and then introduces DNA double-strand breaks (DSBs) at the target sites^1–3^. In mammalian cells, DSBs are repaired through multiple DNA repair pathways: (i) non-homologous end joining (NHEJ), (ii) DNA polymerase theta (Pol θ)-mediated end joining (TMEJ) that operates in a microhomology-dependent manner with as few as 2 to 16 nucleotides of homologous sequence^4^, (iii) single-strand annealing (SSA) that uses homologous repeat sequences to bridge DSB ends, (iv) homologous recombination (HR), and (v) homology-directed repair (HDR)^5, 6^. Because NHEJ, TMEJ, and SSA are error-prone, they frequently lead to insertion or deletion or both (collectively referred to as indel) mutations during the repair process^7–9^. In contrast, the HR and HDR pathways are error-free, thus genome editing that uses single-stranded oligodeoxynucleotides (ssODN) or double-stranded donor plasmids flanking homology arms has higher fidelity by leveraging these repair pathways^10, 11^.

Although most deletions generated by CRISPR nucleases were shorter than 20 base pairs (bp), it was observed that more than 20% of the mutations in mouse embryos induced by CRISPR-Cas9 were unintended deletions longer than 250 bp^12^. Moreover, DSBs induced by CRISPR nucleases can induce chromosomal rearrangements, including chromosomal depletion and translocation^13–15^. Because chromosomal translocation involves off-target cleavage sites and the on-target cleavage site, high-fidelity Cas9 nucleases reduce translocation rates^16^. Unfortunately, high-fidelity Cas9 nuclease do not overcome the issue of unintended large deletions that occur at on-target sites. Another variation of the CRISPR-Cas9 system involves using base editors (BEs) and prime editors (PEs), which employ partially inactivated Cas9 (nCas9) that generates single-strand nicks^17–19^. Whether these nCas9-based genome editing systems also generate large deletion is not clearly established^20–24^. Identifying the repair pathways that produce large deletions following genome-editing tools, with Cas9 or nCas9, will not only provide key insight into the biology of this process but enable the development of strategies to mitigate large deletion events.

Determining the frequency of deletion events >100 bp is challenging using short-range high-throughput sequencing methods. Instead, such events are detected with either a long-range amplicon sequencing method using short-read sequencers, such as Illumina platforms, or a long-read sequencing method, using Pacific Biosciences (PacBio) or Oxford Nanopore Technologies (ONT) platforms^25^. In this study, we modified a long-range amplicon sequencing method and developed a novel k-mer sequence alignment algorithm, and we used this strategy to accurately determine CRISPR-induced large deletions. We observed that large deletions were mostly accompanied by small indels in human cells, including cancer-like cells (HeLa, HEK293T, U2OS, K562), fibroblasts, primary T cells, and human embryo stem cells (H9). We then identified large deletion-associated repair pathways through CRISPR interference (CRISPRi) screening experiments with ONT Nanopore sequencing, establishing that TMEJ is the dominant pathway generating large deletions. We found that BEs generate large deletions through the base excision repair (BER) pathway, especially in the presence of an apurinic/apyrimidinic site (AP site) and that PEs particularly generate large deletions in the presence of additional nicking guide RNAs (ngRNAs). Our findings reveal mechanistic challenges in using genome-editing tools and provide insight for avoiding these issues in research and therapeutic applications.

## Results

### Optimizing a long-range amplicon sequencing method to determine small indels and large deletions with a high accuracy

Neither the PacBio Single-molecule real-time (SMRT) sequencing nor ONT Nanopore sequencing platforms for long-read sequencing have sufficient sequencing accuracy (PacBio ∼10% and ONT 2%-15%)^26^ to distinguish 1-bp deletion, 1-bp insertion, and substitution events. Furthermore, these methods have a length bias: shorter DNA fragments are read more easily and thereby are overestimated compared to longer DNA fragments (**Supplementary** Fig. 1)^27^. Thus, currently available long-read sequencing platforms lack sufficient accuracy to identify both large deletions and small indels. Our first task was to establish a precise sequencing method to simultaneously determine small indels and large deletions with a high accuracy.

The Illumina platform has the highest sequencing accuracy (>99.2%)^28^. Therefore, we optimized long-range amplicon sequencing using Illumina sequencers combined with long-range polymerase chain reaction (PCR) and DNA fragmentation (**Figure 1a**). The four steps in our sequencing were as follows: (i) extract genomic DNA (gDNA) from CRISPR-Cas9-treated and non-treated cell lines, (ii) amplify ∼10 – 15 kb of DNA involving the CRISPR target sites using PCR optimized for both gDNAs, (iii) fragment the amplified products into ∼300 bp and prepare next generation sequencing (NGS) libraries through end-repair, dA-tailing, adaptor ligation, and PCR enrichment, (iv) acquire the sequencing data with an Illumina Miniseq. To quantify large deletions, small deletions, and small insertions in the bulk cell populations, we used a novel k-mer alignment algorithm (**Supplementary** Fig. 2).

**Figure 1.**
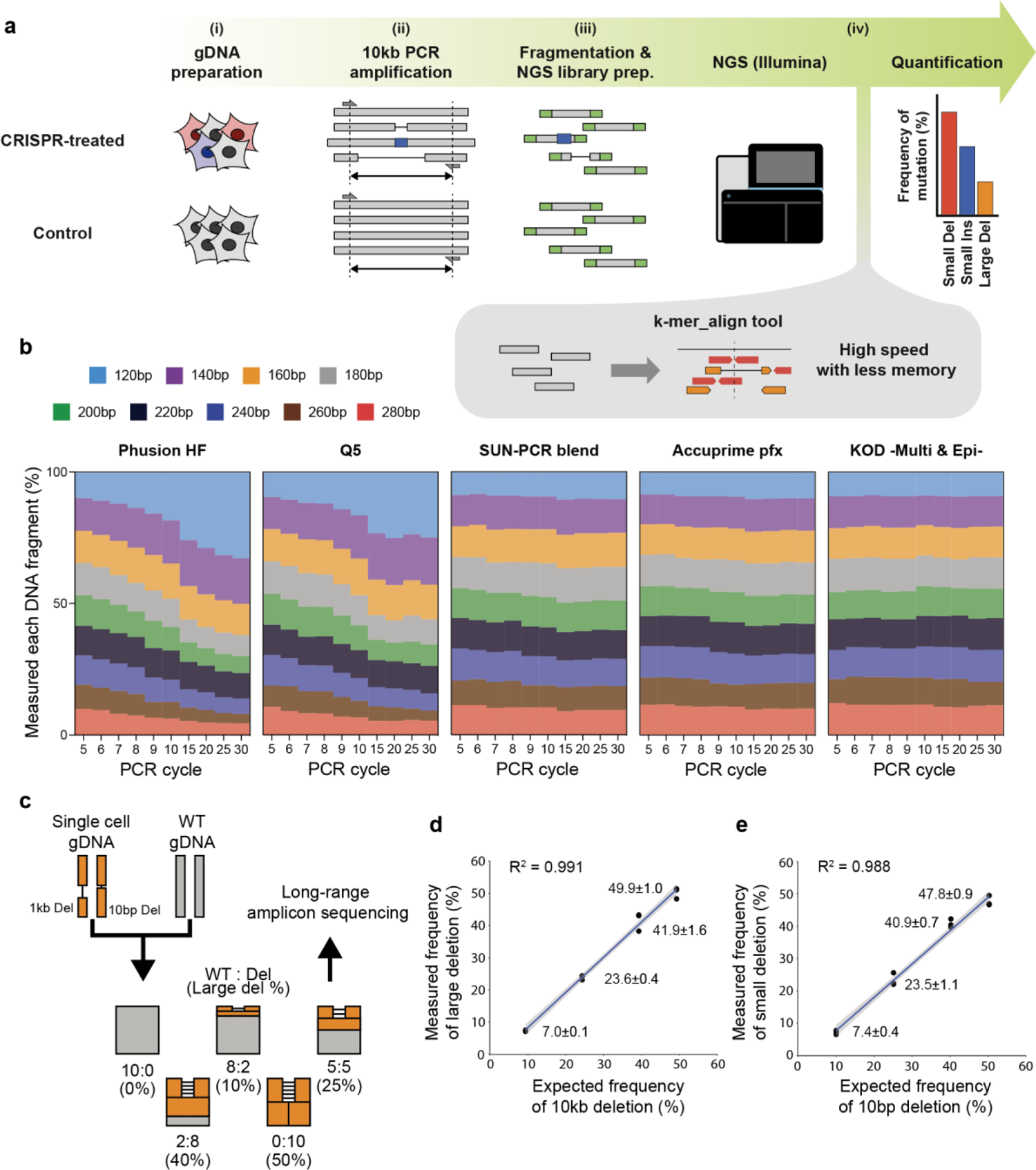
Scheme of long-range deep sequencing and proof of concept. **a**, Schematic of long-range amplicon sequencing. DNA sequences (10 kb) are amplified from genomic DNA isolated from CRISPR-treated and non-treated cells. After fragmentation to ∼300 bp, the fragmented DNA is sequenced by Illumina. The resulting long-range amplicon sequencing data are analyzed with our k-mer alignment program and the frequencies of small mutations and large deletions are calculated. **b**, Comparison of length bias according to type of DNA polymerase and PCR cycle. Nine sizes of DNA fragments were mixed and sequenced with Illumina under various PCR cycle conditions. **c**, Schematic of validation experiments for the long-range amplicon sequencing workflow. Wildtype (WT) genomic DNA (gray) and genomic DNA (orange) from a cell line with a 10 bp and 1 kb deletion in *ST3GAL4* were mixed in a ratio of 10:0, 8:2, 5:5, 2:8, and 0:10. **d-e,** Comparing the expected mutation frequency and measured mutation frequency for large deletions (**d**) and small deletions (**e**).

In this protocol, PCR-based DNA amplification of the long DNA region (∼10 – 15 kb) is a critical step. To mitigate the length bias of DNA polymerases, we tested five DNA polymerases: Phusion HF, Q5 that is used in DNA library prep kit for Illumina, Accuprime pfx^29^, KOD-Multi & Epi-DNA polymerase that is a modified KOD DNA polymerase to reduce amplification bias during PCR by addition of the elongation accelerator^30, 31^, and SUN-PCR blend^32–34^. To evaluate the PCR length bias, we prepared nine synthesized DNA fragments in increments of 20 bp from 120 bp to 280 bp with a common forward/reverse primer pair, mixed the fragments in equimolar amounts, and conducted PCR experiments for the mixture with each polymerase (**Supplementary** Fig. 3). KOD-Multi & Epi-DNA polymerase had the least length bias of the 5 polymerases tested (**Figure 1b**). Therefore, we used this polymerase for the long-range PCR amplification step. Another complicated step in the workflow is sequence alignment of short-read NGS data. Pre-existing tools, such as a burrows-wheeler aligner (BWA)-mem, were developed to rapidly map DNA sequence reads to whole genome sequences. Our workflow requires querying sequences that are limited to the PCR-amplified region to detect indels near the CRISPR-mediated cleavage sites. This is a more limited application that does not require a heuristic approach. Thus, we developed a dedicated k-mer alignment algorithm that is compact and is potentially available as an online tool for analysis of CRISPR-treated samples (**Supplementary** Fig. 4).

To confirm the functionality of our optimized long-range amplicon sequencing workflow, we prepared and applied the workflow to two cell lines: One has two wild-type *ST3GAL4* alleles and the other has a 10-bp or a 1,075-bp deletion in each allele (**Figure 1c**). After mixing gDNAs from the two cell lines at ratios of 0%, 20%, 50%, 80%, and 100%, the mixtures were prepared and subjected to the optimized long-range amplicon sequencing. The optimized long-range amplicon sequencing was highly accurate in detecting both the 10-bp and 1,075-bp deletion events (**Figures 1d** and **1e**), showing that our method is an effective tool for measuring large deletions and small indels simultaneously.

### Determination of CRISPR-mediated small indels and large deletions in various human cell lines

Based on the optimized long-range amplicon sequencing, we quantified large deletions and small indels induced by CRISPR-Cas9 nuclease in various human cell lines. We evaluated four cancer or transformed cell lines (HeLa, HEK293T, U2OS, K562), human fibroblasts, and a human embryo stem cell line (H9) (**Figures 2a-2f**). For HeLa and HEK293T cells, we analyzed the frequency of CRISPR-Cas9-mediated deletion or insertion events for one target site each in *HPRT1* and *ST3GAL4* and five target sites in the *WISP3* gene. We defined a small insertion and a small deletion event as shorter than 100 bp, and a large deletion event as larger than 100 bp. Across these seven target sites, the average frequency of small indel events were 37.5 ± 13.2% in HeLa cells and 26.4 ± 11.7% in HEK293T cells, whereas the average frequency of large deletion event was 6.4 ± 4.4% in HeLa cells and 4.42 ± 3.6% in HEK293T cells (**Figures 2a** and **2b**). Small indels were the most common CRISPR-Cas9-mediated events in all of the cells with an average frequency of 17.8 ± 7.8 % in U2OS cells, 18.4 ± 10.1% in K562 cells, 10.6 ± 8.5% in fibroblast cells, and 49.3 ± 21.7% in H9 cells. CRISPR-Cas9-mediated large deletion events were less common with an average frequency of 3.3 ± 1.8% in U2OS cells, 2.4 ± 1.5% in K562 cells, 1.5 ± 1.7% in fibroblast cells, and 2.6 ± 1.6% in H9 cells (**Figures 2c-2f**). The frequency of CRISPR-Cas9-mediated large deletions across all target sites and all six cell lines ranged from 0.2% to 17.5%.

**Figure 2.**
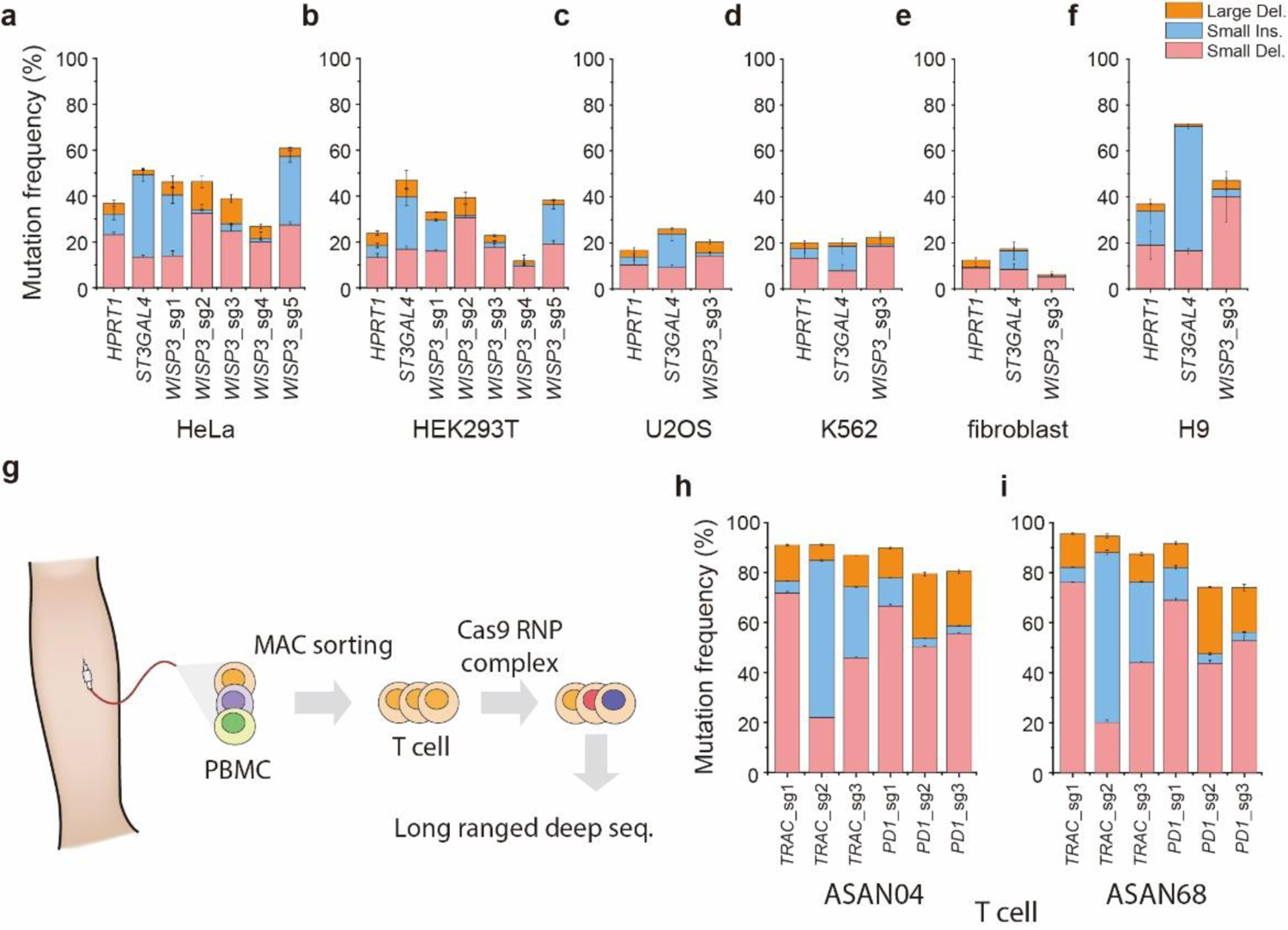
Mutation frequency in various cell lines and human primary T cells. **a-f**, The frequencies of small deletion, small insertion, and large deletion events in HeLa (**a)**, HEK293T (**b)**, U2OS (**c**), K562 (**d**), normal fibroblast (**e**), stem cells (H9) (**f**) after CRISPR-Cas9 gene editing at the indicated genes as determined with long-range amplicon sequencing (n=3). **g,** Schematic of isolation and long-rangd deep sequencing for primary human T cells. **h** and **i**, The frequencies of small deletion, small insertion, and large deletion events in human T cells from two healthy donors, ASAN04 (**h**) and ASAN68 (**i**) subjected to CRISPR-Cas9 editing at the indicated genes as determined with long-range amplicon sequencing (n=3). Error bars, mean±SD.

Because CRISPR-Cas9-based genome editing has been widely adopted in engineering T cells for cancer immunotherapy (clinicaltrials.gov: NCT03399448, NCT03081715, NCT03398967, NCT02793856, NCT03044743), we evaluated CRISPR-Cas9-mediated mutation patterns in human primary T cells. We chose sgRNAs targeting *PD1* and *TRAC1* loci, because these genes are frequently targeted in T cells to enhance T-cell activity^35^. Human T cells were isolated from the peripheral blood mononuclear cells (PBMCs) of two healthy donors (ASAN-04 and ASAN-68) using a magnetic-activated cell sorting (MACS) system and then stimulated by anti-CD3/CD28 beads before introduction of Cas9-sgRNA ribonucleoprotein (RNP) complexes by electroporation (**Figure 2g**). The optimized long-range amplicon sequencing revealed that the average frequencies of small indels were 70.99 ± 11.37% in ASAN04 and 72.03 ±15.46 % in ASAN68, whereas the average frequencies of large deletion events were 15.46 ± 6.68% in ASAN04 and 14.24 ± 6.82% in ASAN68 (**Figures 2h** and **2i**), similar to the previous results^36, 37^. The average frequency of large deletion events in human primary T cells was higher than the average frequency of these events in the cell lines tested.

Using all of the data, we observed that the large deletion frequencies positively correlated with the small deletion frequencies (Pearson corr. = 0.72) and were negatively correlated with small insertion frequencies (Pearson corr. = −0.15), resulting in a low correlation with small indels (Pearson corr. = 0.47) (**Supplementary** Fig. 5).

### Determination by CRISPRi screening that end resection and TMEJ processes play positive roles for generating large deletions

To examine the underlying mechanism of CRISPR-driven large DNA deletion events, we performed a CRISPR interference (CRISPRi) screen targeting 794 genes that were those referenced in Repair-seq^38^ plus genes identified as associated with DNA repair in the Human Protein Atlas website (**Figure 3a**). We introduced a plasmid library with three different sgRNAs per gene and the puromycin resistance gene into an engineered HeLa cell line that stably expresses deactivated Cas9 (dCas9) fused with Krüppel associated box (KRAB) domain to produce gene knockdown^39^. DSBs in the puromycin resistance gene were generated by addition of the Cas9 RNP complex, gDNA was prepared, and ∼5.6 kbp of DNA involving the sgRNA sites or the DSB (puromycin resistance gene) site were amplified and sequenced with the Nanopore MinION. Because we only needed to assess qualitatively large deletion events, we used Nanopore-seq to read long range region. However, to account for the length-dependent bias in Nanopore-seq (**Supplementary** Fig. 1), each sequencing read was adjusted according to its total length and then classified as wild-type, insertion, deletion, and large deletions (those > 100 bp). We compared Nanopore-seq data from cells with or without inhibition of the 794 target genes (**Supplementary** Fig. 6 and **Supplementary Table 1**).

**Figure 3.**
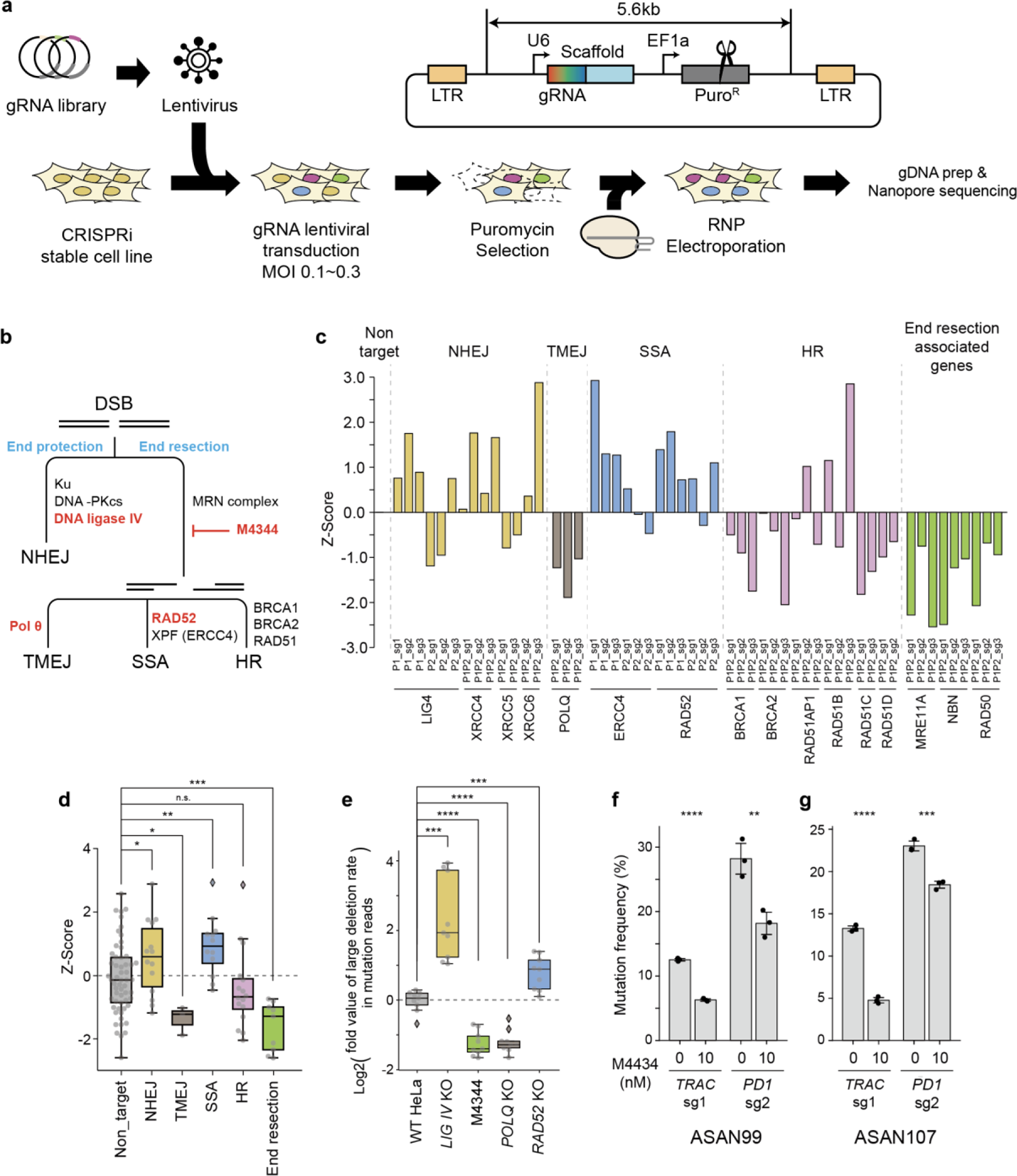
Mutation frequency in knockdown and knockout cell lines. **a**, Schematic of CRISPRi screening with Nanopore-seq. CRISPRi stable cells were infected with a lentiviral library. The infected cells were selected with puromycin and transfected with Cas9 RNP complex targeting the puromycin resistance gene using electroporation. The regions with gRNA or repair outcome (puromycin resistance gene) were amplified and sequenced using Nanopore sequencing. **b**, Model of DNA double-strand break (DSB) repair pathway. Red colored words are genes and chemical (M4344) used in experiments for KO cell lines or pharmacological inhibition of specific DNA repair processes. **c**, Data from CRISPRi screening with Nanopore-seq plotted as Z-scores of the frequency of large deletion events in cells with the indicated gene knocked down. **d**, Z-score comparison of the difference in large deletion frequency between negative control (non-targeting) cells and those with the indicated repair pathway compromised. Z-score was calculated using the mean and distribution of non-targeting sample. P-values were calculated using Mann-Whitney tests. **e**, Box plot graph to compare large deletion rate in wild-type and the indicated KO cells subjected to CRISPR-Cas9. **f** and **g**, The frequency of large deletion events in human primary T cells, which was analyzed in two healthy donors ASAN99 (**f**) and ASAN107 (**g**) using long-range amplicon sequencing (n=3; error bar, mean±SD). P-values in **e**-g were calculated using student’s *t*-test (n.s., not significant, * *P* < 0.05, ** *P* < 0.01, *** *P* < 0.001, **** *P* < 0.0001).

DNA repair pathways can be divided into end protection and end resection^40^ (**Figure 3b**). After DSBs occur, the end protection pathway proceeds to NHEJ, which induces blunt-end ligation by Ku-XRCC4-DNA ligase IV complex^40^. In contrast, the end resection pathway proceeds to TMEJ, SSA, or HR, according to the 5’ end resection sequence and the existence of homology sequences in neighboring DNA^40^. Using a previously defined classification scheme, we categorized the genes into those associated with NHEJ (*LIG4, XRCC4, XRCC5,* and *XRCC6*), end resection (*MRE11A, NBN,* and *RAD50*), TMEJ (*POLQ*), SSA (*ERCC4* and *RAD52*), and HR (*BRCA1, BRCA2, RAD51AP1, RAD51B, RAD51C,* and *RAD51D*). Z-scores for large deletions in the puromycin resistance gene region for each of the CRISPRi-targeted genes within each category indicated that fewer large deletions occurred when *POLQ* of TMEJ or genes associated with end resection were knocked down and large deletions were more common when SSA genes were knocked down (**Figure 3c**). Consistent with these findings, we calculated Z-scores of the difference in large deletion frequencies in cells with and without inhibition of target genes and found a significant reduction in large deletions in cells with either reduced activity of TMEJ (P-value=9.63×10^-3^) or the end resection pathway (P-value=7.71×10^-5^) and a significant increase in cells with reduced activity of SSA (P-value=2.19×10^-3^) (**Figure 3d**). Collectively, the screening results indicated that large deletions are primarily generated by TMEJ following the end resection pathway.

### Confirmation by knockout that end resection and TMEJ processes generate large deletions

To confirm the function of key genes in generating large deletions, we constructed individual knock-out (KO) cell lines using CRISPR-Cas9 or disrupted specific repair pathways pharmacologically. We used the following strategies to impair specific repair pathways in HeLa cells: M4344 to pharmacologically inhibit ATR activity and suppress end resection, *Ligase IV* (*LIG4*) KO to suppress NHEJ, *POLQ* KO to suppress TMEJ, and *RAD52* KO to suppress SSA (**Figure 3b**). We analyzed the mutation pattern in each cell line or condition for three CRISPR-Cas9 targeted genes (*HPRT1, ST3GAL4, WISP3*). Consistent with the CRISPRi screen, we found that *POLQ* KO (P-value=1.46×10^-6^) or the addition of M4344 (P-value=8.83×10^-7^) significantly reduced large deletion rates, whereas *RAD52* KO (P-value=5.64×10^-4^) increased the large deletion rates. In contrast to variable effect of *LIG4* knockdown in the CRISPRi screen, the *LIG4* KO cell line showed a significant increase in large deletion frequency (P-value=3.06×10^-4^), suggesting that blocking either the end protection process or the subsequent NHEJ can enhance the occurrence of large deletions. Both the CRISPRi and KO experiments indicated that the end resection process and the following TMEJ play major roles in generating large deletions after CRISPR-mediated gene editing.

### A strategy to reduce large deletions in primary T cells

As a potential application in T cell engineering, we examined whether M4434 reduced large deletion frequencies in human primary T cells. After introducing CRISPR-Cas9 by electroporation into human T cells, we exposed the cells to various concentrations of M4434 (0 nM, 1 nM, 5 nM, 10 nM, 25 nM, and 50 nM). Cell viability and number were substantially decreased when the M4344 concentration was over 25 nM, and the frequency of large deletions was reduced to similar amounts at M4344 concentrations from 10 nM to 50 nM (**Supplementary** Fig. 7). Therefore, we selected 10 nM as the optimal concentration M4434 to assess the effect on large deletion events in human T cells.

We analyzed the effect of 10 nM M4344 on the CRISPR-Cas9-induced mutation pattern on *TRAC* and *PD1* in human T cells acquired from two healthy donors (ASAN99 and ASAN107). For this analysis we used the optimized long-range amplicon sequencing method. M4344 decreased the frequency of large deletion events between 35% and 80%, and the reductions were significant: *TRAC* P-value=5.41×10^-6^ and *PD1* P-value=8.52×10^-3^ in ASAN99, *TRAC* P-value=1.14×10^-5^ and *PD1* P-value=8.03×10^-4^ in ASAN99 (**Figures 3f**, **3g** and **Supplementary** Figs. 7d, 7e). TMEJ-mediated DNA repair is mediated by microhomology, meaning that repair can occur with as few as 2-16 nucleotides of homologous sequence^4^. We found that the microhomology-mediated deletion frequencies were also decreased (**Supplementary** Fig. 8), indicating that M4344 prevents the large deletion events generated by TMEJ. Collectively, these data support evaluation of M4344 or drugs that reduce end resection or TMEJ pathways as co-effectors with CRISPR-Cas9-mediated genetic engineering in therapeutic applications to mitigate unintended large deletions.

### Generation of large DNA deletions by cytosine base editors and adenine base editors

In contrast with Cas9 nucleases, BEs employ nCas9 containing the D10A mutation, which is impaired in the ability to generate DNA DSBs. Instead of introducing DSBs at target sites, BEs modify single nucleotides in the DNA sequence. BEs, including cytosine base editor (CBE) and adenine base editor (ABE), introduce infrequent indels^17, 18, 41^. Thus, it is possible for BEs to generate large deletions. It was reported that CBE variants without a uracil glycosylase inhibitor (UGI) have relatively high indel frequencies^41, 42^, thus we hypothesized that BE platforms without UGI induce large deletions with a high frequency. The role of the UGI is to inhibit cellular uracil DNA glycosylase (UNG), the enzyme that excises uracil^43, 44^. After uracil excision, cellular DNA AP lyase introduces a nick on the non-target strand of the sgRNA, leading to BER^45^. Therefore, we predicted that UGI-lacking BEs generate DSBs and consequently large deletions during the BER process through a nick on each strand— one by nCas9 (D10A) and one by AP lyase. ABE variants also induce cytosine editing in preferred motifs, suggesting the possibility of ABE introducing large deletions during bystander cytosine editing^42, 46^.

We measured large deletion frequencies with six different BE platforms: a canonical CBE containing UGI^47^, a CBE variant without UGI [named CBE (ΔUGI)], a CBE variant without UGI and with additional UNG (named CGBE1)^41^, a canonical ABE^17^, an ABE variant with UGI (named ABE-UGI), and an ABE variant containing engineered hypoxanthine excision protein N-methylpurine DNA glycosylase (MPG) (named AYBE)^48^ (**Figure 4a**). We introduced each BE platform, or inactivated Cas9 (dCas9) as a control, into HEK293T cells and targeted four endogenous targets (*ABLIM3, FANCF, KLHL29, BRD4*); then we conducted optimized long-range amplicon sequencing to evaluate large deletion frequencies.

**Figure 4.**
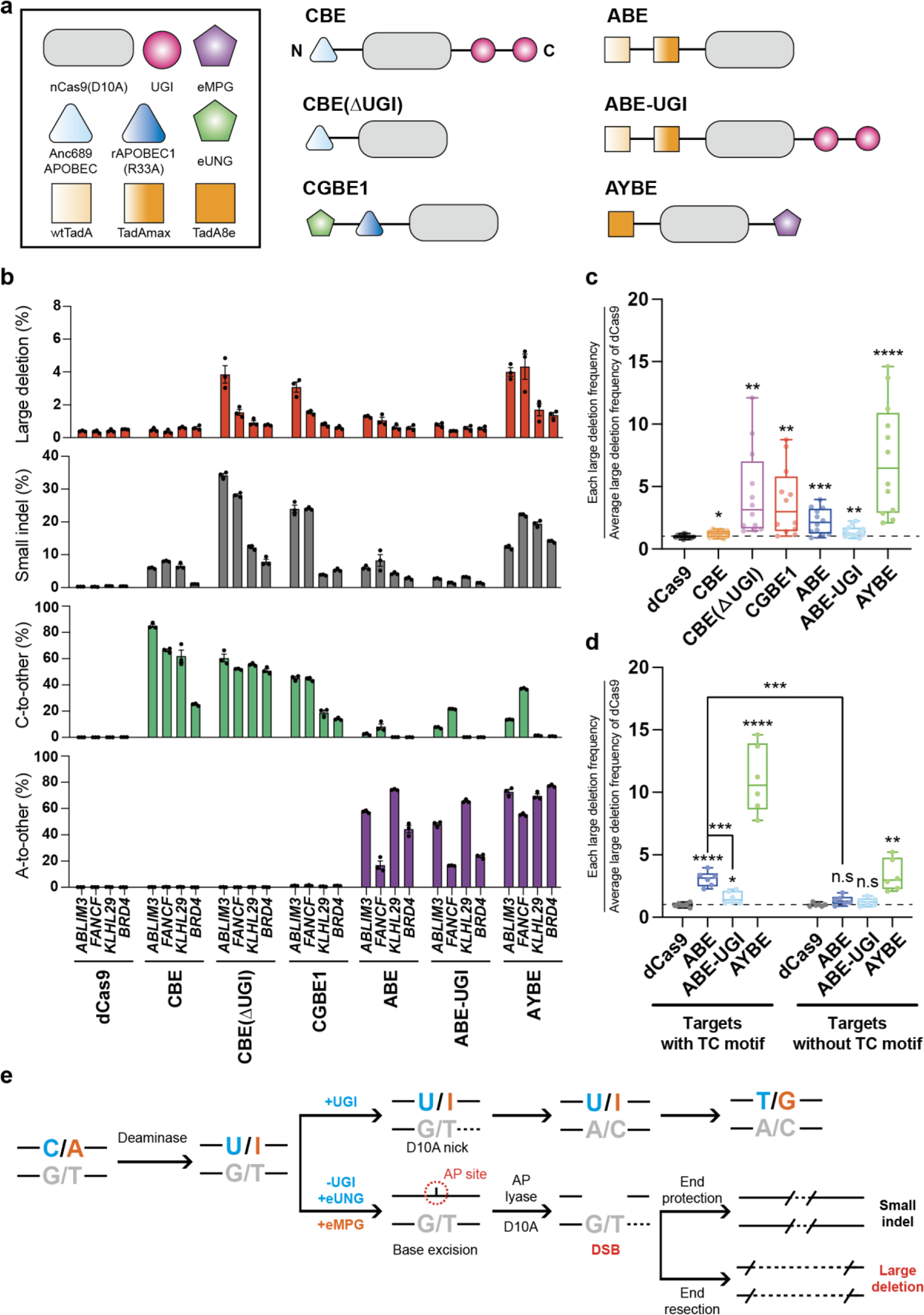
DNA large deletions are generated by base editors. **a.** Schematic diagrams of six different BEs. N and C indicate the amino-terminal and carboxy-terminal ends, respectively. The version of CBE and ABE is AncBE4max and ABEmax, respectively. **b.** Editing outcomes of dCas9 (inactivated Cas9) and the indicated BEs targeting *ABLIM3*, *FANCF*, *KLHL29*, and *BRD4* (*n*=3, mean ± SEM). **c.** Summarized box plot graph to compare the large deletion frequency among the BEs targeting *ABLIM3*, *FANCF*, *KLHL29* and *BRD4*. *P* = 0.0249 for CBE, *P* = 0.0028 for CBE(ΔUGI), *P* = 0.0020 for CGBE1, *P* = 0.0005 for ABE, *P* = 0.0062 for ABE-UGI, and *P* < 0.0001 for AYBE compared to the dCas9 by student’s *t*-test. **d.** Analysis of the large deletion frequency of ABE with or without UGI at targets containing a TC-motif (TC target, *ABLIM3* and *FANCF*) and targets without a TC-motif (Non-TC target, *KLHL29* and *BRD4*). Statistical analysis was performed by student’s t test. Within the TC target, *P* < 0.0001 for ABE, *P* = 0.0260 for ABE-UGI and *P* < 0.0001 for AYBE compared to the dCas9. *P* = 0.0007 for ABE comparing with ABE-UGI at the TC target. Within the non-TC target, *P* = 0.0564 for ABE, *P* = 0.1264 for ABE-UGI and *P* = 0.0010 for AYBE compared to the dCas9. *P* = 0.0001 for ABE at TC target compared to the ABE at non-TC target. **e.** Schematic of potential cellular mechanisms generating DNA large deletion by BEs. Uracil or inosine excision by endogenous uracil DNA glycosylase (UNG) or engineered protein N-methylpurine DNA glycosylase (eMPG), nicking of the non-target strand by nickase Cas9(D10A), AP site generation by DNA Apurinic or apyrimidinic site lyase (AP lyase), followed by DNA double strand break (DSB) formation. \**P* < 0.05, \*\**P* < 0.01, \*\*\**P* < 0.001 and \*\*\*\**P* < 0.0001. The description of n.s means non-significant.

Each BE platform generated large deletion mutations; however, the frequency varied both for each platform and each targeted gene (**Figure 4b**). Compared to the canonical CBE, which had a maximum frequency of large deletion events of 0.8%, CBE (ΔUGI) and CGBE1 generated higher frequencies of large deletion events with maximum frequencies of 4.8% and 3.5%, respectively. The canonical ABE had a large deletion frequency maximum of 1.42%. In Figure 4c, each large deletion frequency was divided by the average large deletion frequency with. Overall, introduction of a UGI slightly reduced the frequency and introduction of the eMPG increased the frequency substantially with a range of 4.41% to 1.21% and 2.21% to 7.19%, respectively (**Figure 4c**).

For ABE platforms, we observed that target sequences including a TC motif within the editing window, as in *ABLIM3* and *FANCF*, had larger frequencies of both small indels and large deletions compared to target sequences without a TC motif, as in *KLHL29* and *BRD4* (**Figures 4b** and **4d**), indicating that bystander C-to-U conversion by ABE and the resulting AP site produces DSBs and subsequent large deletion events. Both small indels and large deletion events occurred at lower frequencies at targets with TC motifs with ABE-UGI, consistent with the UGI inhibiting the production of DSBs produced through a mechanism similar to that observed for the CBEs (**Figure 4d**). Compared to targets with a TC motif, those without a TC motif had a lower frequency of either large deletion events or small indels due to AYBE (**Figures 4b** and **4d**). However, the frequency of large deletions due to AYBE at the target sites without TC motif was substantially higher than that of ABE. Thus, inosine excision by eMPG of AYBE likely enables DSB generation and large deletions through a mechanism like that with the uracil excision by UNG in which Cas9 (D10A) nickase nicks one strand and the BER process nicks the other strand (**Figure 4e**).

### Generation of large deletion events by prime editors

In contrast to BEs, PEs employ nCas9 containing H840A mutation and there are two representative PE platforms, PE2 and PE3^19^. PE2 introduces a single nick on the non-target strand of the pegRNA, whereas PE3 introduces a double nick on each strand of the DNA through the addition of a nicking guide RNA (ngRNA)^19^. Empirically, we and other groups observed that PEs are often accompanied with unwanted small indels^19, 49^. In addition, compared with nCas9 (D10A), the cleavage activity of nCas9 (H840A) is not completely impaired and introduces DSBs with a low efficiency^50^. The potential for DSBs that result in large deletions is particularly high for PE3^21, 51^.

We measured large deletion frequencies induced by PE2 or PE3 at substitution targets in *HEK3* and *HEK4*, deletion targets in *FANCF* and *HEK3*, and insertion targets in *RNF2* and *HEK3*. As expected, PE2 had a lower frequency of large deletion (maximum 1.2%) than PE3 (maximum 24.3%) (**Figures 5a-5c**). Similar to the BE platforms, the frequency of PE-induced large deletion events varied by target (**Figure 5a**). Particularly with PE3, even in the same target, large deletion frequencies varied according to the targeted edit: Compare *HEK3* 1-10del and *HEK3* 1CTTins, each with +90 ngRNA and *HEK4* +2GtoT with the ngRNA +52 or +74 (**Figure 5a**). These results suggested that the sequence in proximity to the prime editing site might contribute to the potential for large deletion events.

**Figure 5.**
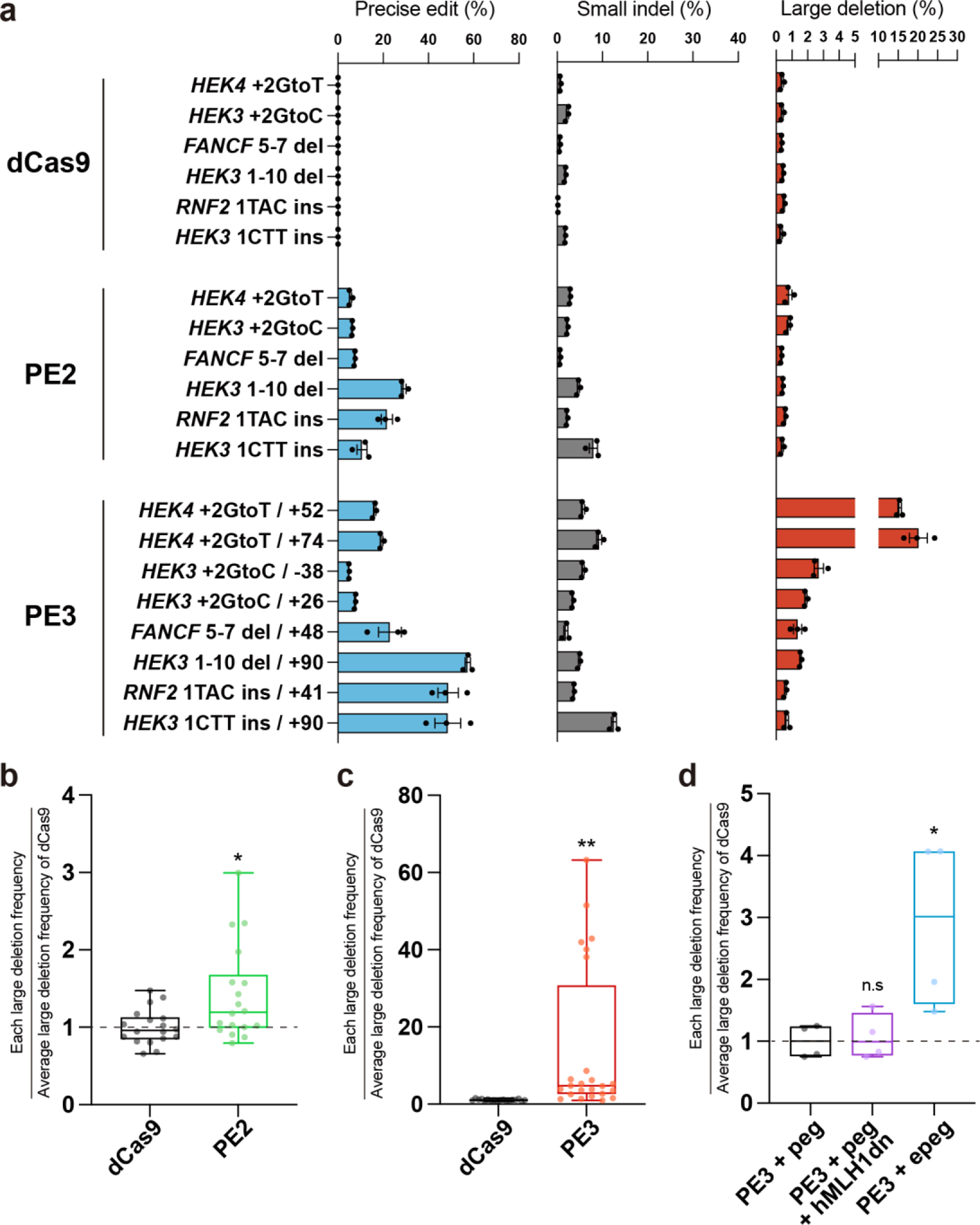
Prime editors can generate DNA large deletion especially with the additional nicking gRNA. **a.** Editing outcomes of dCas9, PE2, and PE3 with substitution, deletion, or insertion edits. Two targets were tested for each type of editing: *HEK4* and *HEK3* for substitution, *FANCF* and *HEK3* for deletion, *RNF2* and *HEK3* for insertion. For PE3 strategy, two ngRNAs were tested individually. (*n*=3, mean ± SEM). **b** and **c.** Summarized box plot graph describing the large deletion frequency of PE2 (b) or PE3 (c). *P* = 0.0103 for PE2 and *P* = 0.0056 for PE3 (Student’s *t*-test, two-sided). **d.** The effect of hMLH1dn or epegRNA on DNA large deletions induced by the PE3 system. Analysis was performed with targets +2GtoC with −38 ngRNA in *HEK3*, 5-7del with +48 ngRNA in *FANCF*. *P* = 0.7507 for PE3 + hMLH1dn and *P* = 0.0347 for PE3 + epegRNA compared to the original PE3. \**P* < 0.05, \*\**P* < 0.01; n.s means non-significant.

We investigated whether the mismatch repair (MMR) pathway or increasing the lifetime of the pegRNA affects the formation of large deletions. To determine the effect of MMR, we introduced PE3 with pegRNA and a MMR inhibitory factor, hMLH1dn^52^. Inhibition of the MMR pathway did not significantly affect the frequency of large deletion events compared with those induced by PE3 with pegRNA (**Figure 5d**), suggesting that the MMR pathway has a low contribution for the occurrence of PE3-mediated large deletions. Using an established method to increase pegRNA stability, we introduced PE3 with the engineered pegRNA (epegRNA) and found that this resulted in an increase in large deletion frequencies^53^ (**Figure 5d**). We also compared the effect of epegRNA combined with two enhanced PE systems, PE4max and PE5max^52^, on large deletion frequencies induced during insertion of 1 TAC into *RNF2* or of 1 CTT into *HEK3*, which were sites with relatively low frequencies of large deletion in the PE2 and PE3 systems with pegRNA (**Figure 5a**). Compared with the occurrence of large deletion events in cells with dCas9, PE4max with epegRNA did not significantly affect the frequency of large deletion events but PE5max with epegRNA induced a significantly greater frequency of large deletion events (P-value=0.0071) (**Supplementary** Fig. 9).

## Discussion

Here, we refined a long-range amplicon sequencing method to enable simultaneous analysis of both large deletions and small indels with high accuracy. To develop this long-range amplicon sequencing method, we determined the PCR polymerase with minimal length bias, optimized the PCR protocol, and developed a dedicated k-mer sequence-alignment algorithm along with an optimized analysis program to eliminate false-positive large deletion reads (**Supplementary** Fig. 10). With this method, we detected and quantified large deletions and small indels in all human cells that we tested. Many groups analyze CRISPR-mediated gene editing outcomes within a short range (<500 bp) of the target site, which can miss large deletion events. We experienced mischaracterization of a CRISPR nuclease-mediated knockin cell line (**Supplementary** Fig. 11). Based on PCR amplification and short-range deep sequencing data, we observed only one pattern for a *EXD2* Halo tag knockin cell line, suggesting that the cell line had homologous knockin at both alleles However, with long-range amplicon sequencing, we found that one allele had a large deletion (740 bp). Thus, this is a heterologous knock-in line. A similar phenomenon was observed with wheat genetically engineered for pesticide resistance in which a large deletion altered the epigenetic landscape and changed the growth properties of the plant^54^. Hence, it is necessary to test for the presence of large deletions after gene editing with CRISPR nucleases.

Reducing unintended chromosomal changes in therapeutic applications, such as in generation of CAR T cells, is critical. Tsuchida et al. demonstrated that large deletions and chromosomal truncations can be reduced by altering the step of T cell activation in experimental protocols^55^. We determined that TMEJ serves as a primary repair pathway for generating large deletions. Understanding the mechanism responsible for large deletion events enables development of mitigation strategies. Indeed, we showed that inhibiting the end resection pathway and the subsequent TMEJ pharmacologically with M4344 reduced CRISPR-Cas9-induced large deletion frequencies in human primary T cells.

BEs and PEs that use Cas9 nickase have the potential to generate DSBs at target sites^21, 50, 51^, but some studies report that BEs and PEs do not generate large deletions^20, 22^. However, our data showed that both BEs and PEs generate large deletions, especially through BER pathway-dependent base editing that creates an AP site for BEs. PE systems that use ngRNA or the stabilized epegRNA showed particularly high frequencies of large deletion.

Gene editing tools based on Cas9 or nCas9 exhibited frequencies of large deletion events that varied by target site and for the nCas9-based systems by type of genetic modification. The first-in-kind CRISPR gene editing drug, named Exa-cel or CASGEVY, was approved in UK and USA in 2023. This is remarkably fast for a therapeutic based on CRISPR-Cas nucleases and begins a new era for gene editing therapy. Our findings showed that the present genome-editing tools need additional validation to ensure large deletion events are not present or these tools need additional engineering to prevent the generation of DSBs and large deletions. Whether there are additional shortcomings yet to be discovered for the clinical application of these genome editing tools remains an open question. The workflow that we developed provides a mechanism to evaluate unintended large or small changes to DNA arising from the application of these gene editing tools.

## ACKNOWLEDGEMENTS

Most analysis of sequencing data was carried out using the computing server at the Genomic Medicine Institute Research Service Center. This research was supported by grants from the National Research Foundation of Korea (NRF) no. 2021R1A2C3012908, no. 2021M3A9H3015389 to S.B. The authors thank Nancy R. Gough (BioSerendipity, LLC) for editorial service.

## AUTHOR CONTRIBUTIONS

S.B. and G.H. conceived this project; G.H., S.-H.L., and H.S.K., performed and analyzed screening experiments; M.O. and G.H. developed bioinformatics algorithms; G.H., S.-H.L., and O.H. performed cell experiments; S.K. and H.-K.J. performed T-cell experiments; C.H.K., S.K., and S.B. supervised this project; G.H., S.-H.L., and S.B. wrote the manuscript with the help of all other authors.

## Additional information

Supplementary Information accompanying this paper is available at http://

## Materials and methods

### Generation of sgRNA-encoding plasmids

Target sequences were designed using Cas-Designer^56^ to avoid off-target sequence (up to 2 mistmaches). The list of oligomers for target sequence is in **Supplementary Table 1**. pRG2 GG expression vector was digested using *Bsa1* restriction enzyme. sgRNA oligos, with overhangs complementary to the digested vector, were ordered from Macrogen (Korea) and Cosmogenetech (Korea). These oligos— comprising both the upper and lower strands—were then annealed to produce double-stranded oligo deoxynucleotides (dsODN). The annealed dsODNs were ligated into the digested expression vector using T4 DNA ligase (Enzynomics) and incubated for 1h at room temperature. The ligation mixture was transformed into DH5a competent cells using the heat-shock method and cultured overnight at 37°C. Individual colonies were selected and grown in LB media for 16 hours at 37°C in a shaking incubator. The plasmids were isolated using Exprep™ Plasmid SV kit (GeneAll).

### Cell culture and transfection for cancer cell lines and fibroblasts

HeLa (ATCC®, CCL-2™), HEK293T (ATCC®, CCL-3216™) cells, and U2OS (ATCC HTB-96) cells were maintained in Dulbecco’s Modified Eagle Medium (DMEM) supplemented with 10% fetal bovine serum (FBS), 100 unit/mL penicillin, and 100 unit/mL streptomycin. K562 (ATCC CCL-243) cells were maintained in Roswell Park Memorial Institute (RPMI) 1640 Medium supplemented with 10% fetal bovine serum (FBS), 100 unit/mL penicillin, and 100 unit/mL streptomycin. Normal fibroblasts [ThermoFisher, Human Dermal Fibroblasts (C0045C)] were maintained in Dulbecco’s Modified Eagle Medium (DMEM) supplemented with 20% fetal bovine serum (FBS), 100 unit/mL penicillin, and 100 unit/mL streptomycin.

HeLa and HEK293T were transfected with Lipofectamin 2000 (Invitrogen). Before transfection, 1 × 10^5^ cells from each well were seeded in 24-well plates. SpCas9 expression plasmids (750 ng) and sgRNA expression plasmids (250 ng) were mixed with 100 µl of Opti-MEM medium and 2 µl of Lipofectamin 2000 and incubated for 20 min at room temperature. The prepared mixture was added to the seeded wells. After 24 hours, the culture media were replaced with fresh media. U2OS and K562 were transfected with Neon transfection system (Invitrogen). cells (2.5 × 10^5^) were transfected with 750 ng of Cas9 expression plasmids (750 ng) and sgRNA expression plasmids (250 ng) with the following parameters: 1050 V, 30 ms, 2 pulse for U2OS and 1350 V, 10 ms, 4 pulse for K562. Normal fibroblasts were transfected with Amaxa P3 primary cell 4D-nucleofector kit using program DS-137. All cells were analyzed 3 days after transfection.

### Cell culture and transfection for H9 cells

H9 human embryonic stem cells were maintained in Essential 8 (E8) medium (Gibco A1517001) on iMatrix-511 (Matrixome, 892 021). The dissociation of H9 cells into clusters for subculturing was facilitated using ReLeSR (Stemcell Tech., 05873). Subsequently, the cells were transferred and replated in E8 medium supplemented with p160-Rho-associated coiled-coil kinase (ROCK) inhibitor Y-27632. Before electroporation, TrypLE (Gibco, 12604013) was used to generate a suspension of single cells. Cells (1 × 10^5^) were electroporated with 250 ng of sgRNA-encoding plasmid and 750 ng of Cas9 expression plasmid using a NEON system (ThermoFisher) at 1050 V for 30 ms (two pulses). Cells were then seeded in 48-well plates in E8 supplemented with Y-27632 (10 μM) for 24 hours. After three days of culturing, gDNA was isolated.

### Generation of Cas9 ribonucleoprtein (RNP) complexes for CRISPRi screening

The puromycin targeting sgRNA was synthesized by *in vitro* transcription using T7 RNA polymerase (NEB) and template oligos, and the sgRNA product was purified using RNeasy Mini Kit (Qiagen). *Streptococcus pyogenes* Cas9 (SpCas9) was ordered from Enzynomics. To generate Cas9 RNP complex, SpCas9 and sgRNA were mixed in a ratio of 1:3 and incubated at room temperature for 30 min. These Cas9 RNP complexes were added to the CRISPRi-stable HeLa cell line after lentiviral transduction and puromycin selection for CRISPRi screening.

### Isolation, culture and editing of human primary T cells

Whole blood samples from healthy donors were taken under a protocol approved by the committee of Asan Medical Center. Peripheral blood mononuclear cells (PBMCs) were isolated from the whole blood samples using SepMate PBMC isolation tubes (STEMCEL). The PBMCs were further processed to isolate human primary T cells using MACS based PAN-T isolation kits (Miltenyi Biotec). RPMI-1640 (Gibco) supplemented with fetal bovine serum (10%, gibco), GlutaMAX (2 mM, gibco), sodium pyruvate (1 mM, gibco), non-essential amino acids (0.1 mM, gibco), beta-mercaptoethanol (55 μM, gibco), HEPES (10 mM, sigma) and penicillin-streptomycin (1%, gibco) was used to culture human primary T cells with IL-2 (300 IU/ml, BMI KOREA).

For gene editing, the human primary T cells were stimulated with Dynabeads human T-Activator CD3/CD28 (Thermo Fischer Scientific) at a cell-to-bead ratio of 1:1 for 48 h. After separating T cells from beads using a magnet, the stimulated T cells were electroporated with Neon transfection system (Thermo Fisher). Briefly, 5 μg of recombinant Cas9 (Enzynomics) and 5 μg of *in vitro*-transcribed sgRNA was incubated at 37°C for 10 min to form Cas9 RNP complex, immediately before electroporation. Assembled CRISPR RNPs were added to 0.5 million of activated human T cells resuspended in T buffer and electroporated with a Neon electroporation device (1400 V, 10 ms, 3 pulse). Electroporated cells were transferred into culture vessels containing culture medium without antibiotics. One day after electroporation, culture medium was changed into fresh medium containing antibiotics and cells were maintained at a concentration of approximately 1 million cells per ml of medium.

### Long-range amplicon sequencing

The PCR primers were designed using Primer3Plus^57^ (https://www.bioinformatics.nl/cgi-bin/primer3plus/primer3plus.cgi) or Primer-BLAST^58^ (https://www.ncbi.nlm.nih.gov/tools/primer-blast). The targeted region (∼8 to 15 kb) in gDNA was amplified using KOD multi & epi DNA polymerase (TOYOBO) according to the manufacturer’s protocol. For sequences that were difficult to amplify, primers were replaced or gDNA was re-extracted. Amplified products (1 µg) were purified using AMPure XP bead-based reagent (Beckman Coulter) with 0.95X. The purified DNA samples were fragmented to ∼300 bp with M220 Focused-ultrasonicator (Covaris) according to the manufacturer’s protocol. The fragmented samples were purified with Expin™ PCR SV kit (GeneAll) and prepared as an NGS library with NEBNext® Ultra™ II DNA Library Prep Kit for Illumina® (NEB). The prepared samples were sequenced with MiniSeq High Output Reagent Kit (300-cycles) using MiniSeq to obtain about ∼400,000 to 500,000 reads. The NGS data FASTQ files from CRISPR-treated and non-treated cells were analyzed using the k-mer alignment program.

### Developing k-mer alignment program

We developed a *k*-mer based algorithm for detecting CRISPR-induced DNA alterations without requiring a supercomputer and that can run on a personal computer. The *k*-mer alignment program is efficient to run with limited computational resources. For the cases investigated here, we used a personal computer with (16 GB) memory and (3.4 GHz / 8 cores) CPU. Our software program accepts as input a 10-kbp reference sequence that includes the cleavage site and the paired-end sequencing data in FASTQ format from both CRISPR-treated and non-treated samples. The output quantifies CRISPR-treated large deletions and small indels and provides read alignment results. The alignment program consists of 3 steps: (i) the short read alignment step, (i) short read classification step, and (iii) removal of false-positive large deletions (**Supplementary** Fig. 4).

The first step is the short read alignment task based on a *k*-mer hash table that is constructed using a reference genome. This hash table is used to obtain positions on the reference genome of (*l-k+1*) *k*-mers for a given read of length *l*. To determine alignment of given input, our program identifies the longest region of consecutive overlapping *k*-mers on the reference sequence through the Longest Increasing Subsequence (LIS) algorithm.

The second step is the short read classification task. Based on the read alignment results, our program categorizes the read pairs into three distinct classes, (i) all read pairs are skewed to the cleavage site, (ii) all read pairs are mapped without splitting and the cleavage site passes between reads, and (iii) one of the reads is split and passes through the cleavage site. Read pairs in category (i) are considered wildtype. Read pairs in categories (ii) and (iii) are considered as potential CRISPR-derived variant candidates and are compiled into a candidate list.

The third step involves the elimination of false-positive large deletions. It is possible for variant candidate read pairs to be present in non-treated samples, primarily due to inherent biases associated with PCR amplification. This phenomenon is not specific to CRISPR-treated samples, but it is a consequence of the characteristics of the reference sequence and affects both CRISPR-treated and non-treated datasets. To reduce the influence of false positives, our program performs a mapping of candidate read pairs to the left and right positions within the reference sequence and identifies clusters of these candidates by implementing *k*-means clustering. The appropriate value for *k* (the number of clusters) is determined using the Bayesian information criterion, which enables the automated selection of the optimal *k* selected based on the characteristics of the dataset. After cluster selection, those clusters that are common to both CRISPR-treated and non-treated datasets are categorized as a result from PCR bias rather than CRISPR-induced variation. These co-occurring clusters are then excluded from the candidate list. Finally, the remaining read pairs in the candidate list are identified as CRISPR-derived variants.

Read pairs with a deletion length of 50 base pairs or more within the curated list of candidate variants are determined as a large deletion. Read pairs with insertions or deletions within a range of ±13 base pairs around the cleavage site are categorized as small indels. To quantify the identified reads, our program enumerates the occurrence of normal mappings, small indels, and large deletions within read pairs passing through the cleavage site (±13 base pairs). Our program then calculates the relative frequencies at which large deletions, small insertions, and small deletions occur. Let *N*_*T*_ be the total number of reads passing the cut site (±13 base pairs), including large deletions. *N*_*L*_ is the number of large deletions. *N*_*S*_, *N*_*D*_ are the number of reads for small insertions and small deletions. The relative frequency of large deletions calculated by *N*_*L*_/*N*_*T*_. Small indels are calculated by *N*_*S*_ + *N*_*D*_ / *N*_*T*_. Deletion analysis was performed with the developed *k*-mer alignment program using a local computer version with an added web front-end.

### CRISPRi library construction

Our custom CRISPRi library was constructed using pAX198 (Addgene #173042) that includes pU6-sgRNA-EF1a-Puro-T2A-BFP. Using the Repair-seq dataset and genes associated with DNA repair in the Human Protein Atlas website, we selected 794 genes associated with DNA repair and categorized them into specific repair processes or pathways. For each gene, three CRISPRi gRNA sequences were sourced from the hCRISPRi-v2.1 library developed by the Weissman group (**Supplementary Table 2**) and 60 non-targeting gRNAs from Repair-seq were added in the CRISPRi library. The oligonucleotide library was procured from GenScript (USA) and subsequently amplified employing the Phusion® High-Fidelity DNA Polymerase (NEB). The amplified product and pAX198 plasmids were digested using FastDigest Bpu1102I and FastDigest BstX1 (ThermoFisher). Desired DNA fragments from the digested pAX198 were isolated through a 1% agarose gel and subsequently purified using the Expin™ Gel SV kit (GeneAll). The digested oligo library was separated and the required DNA sequence was extracted from a 10% PAGE gel. This gel-extracted DNA was then purified using isopropanol precipitation. The digested oligo library and plasmid backbone were ligated using T4 DNA ligase. The ligation mixture was then purified with AMPure XP beads. The ligated product was transformed into MegaX DH10B T1R Electrocomp™ Cells (ThermoFisher) using MicroPulser Electroporator (BioRad). After confirming more than 90,000 colonies, the plasmid library was obtained using NucleoBond Xtra Midi EF kit (Macherey-Nagel). The oligo library was confirmed using nested PCR and Illumina sequencing.

### Lentivirus preparation

Lentivirus was produced in the Lenti-X 23T cell line (Takara Bio). Transfection was carried out with psPAX2 (Addgene #12260), pMD2.G Addgene #12259), and library plasmids using polyethyleneimine (Sigma-Aldrich). The cultured medium was replaced one day post-transfection. Two days later, lentivirus-containing medium was harvested and filtered through a 0.45-µm syringe filter. The lentivirus was concentrated using the Lenti-X concentrator (Takara Bio). The viral titer was determined by performing lentiviral transductions at varying concentrations in a 48-well plate format. After titration, the lentiviral library was aliquoted and stored at −80°C.

### CRISPRi screening with Nanopore sequencing

HeLa CRISPRi cells were generated by lentiviral integration (∼3 to 5 MOI) using the dCas9-KRAB-blast plasmid (Addgene #89567), followed by single cell isolation. Prior to lentiviral transduction, HeLa CRISPRi cells were cultured at a density of 5 × 10^5^ cells in a 100-mm dish. The following day, the sgRNA library lentivirus was added to the cultured HeLa CRISPRi cells in the presence of 8 µg/mL polybrene. After 24 hours, the culture medium was replaced with fresh medium. Another 24 hours later, cell selection was initiated with 2 µg/mL puromycin and continued for 2 to 3 days. Post-selection, the culture medium was replaced with fresh medium and the cells were cultured for 6 to 8 days to allow gene repression to occur. Subsequently, Cas9 RNP complex targeting the transduced puromycin resistance gene was transfected into the sgRNA library transduced CRISPRi stable cell line using Neon transfection system (**Figure 3a**). Three days post-transfection, gDNA was extracted using the NucleoSpin Blood XL, Maxi kit (Macherey-Nagel). Cell culture was performed whenever the cells had grownto 90% of the cell plate.

Half of the extracted gDNA was amplified to generate fragments of ∼5 to 6 kb using the KOD multi & epi DNA polymerase. These amplified fragments were then purified with AMPure XP beads. The purified samples were sequenced on MinION (Oxford Nanopore) using ligation sequencing kit V14 (Oxford Nanopore) and MinION flow cell R10.4.1 (Oxford Nanopore) according to the manufacturer’s protocol. The sequencing process ran at a speed of 260 bps, and base calls were made on the resulting data using guppy (Oxford Nanopore) with super high accuracy mode.

### Analysis for CRISPRi screening with Nanopore sequencing

Fastq files were aligned to the reference genome using the guppy aligner with default settings. To identify gRNA sequences within the sequencing data, we utilized BWA-mem. A gRNA reference FASTA file was constructed by appending 10 bp from the reference sequence to both ends of the gRNA sequence and the gRNA reference FASTA file was indexed using BWA. The gRNA sequence of each read was obtained and aligned with BWA-mem, applying the parameters “-k10-A4-B2-O2”. The sequencing results were saved as files according to gRNA. To ensure data accuracy, we discarded reads where sequences downstream of both the gRNA and the the Blue fluorescent protein (BFP) did not align. If the deletion was more than 100 bp and the deletion spanned a region within 100 bp of the cleavage site, the deletion was classified as a large deletion mutation. Because Nanopore sequencing has a bias depending on the length of the DNA fragment (**Supplementary** Fig. 1), the ratio of the length of the DNA fragment to the reference was used instead of the count so that the ratio would decrease as the length of the deletion became longer. Based on the results of 60 non_targeting gRNAs, the Z-Score of large deletion for each gRNA was calculated.

### Knock-out cell line generation

To generate three different gene (*LIG IV*, *POLQ*, *RAD52*) knock-out HeLa cell line, 750 ng of Cas9 expression plasmid and 250 ng of sgRNA targeting upstream exon of each gene were transfected with 2ml of Lipofectamine 2000 reagent (Invitrogen) into HeLa cells. After 72 hours, CRISPR-treated HeLa cells were distributed as a single cell into each well of 96-well plates. The cell lines were cultured for two weeks, and each genotype was confirmed by an Illumina Miniseq instrument. The Miniseq results were analyzed using Cas-Analyzer (http://www.rgenome.net/cas-analyzer/)^59^. For the complete knock-out, the cell line harboring a frameshift mutation in both alleles was selected.

### M4344 toxicity and effect on CRISPR-induced large deletion events

To inhibit the ATR protein in HeLa cells, we used M4344 (Selleckchem, S9639) according to manufacturer’s protocol. HeLa cells (1 × 10^5^) were seeded in 24-well plates. After 24 hours, the cells were exposed to M4344 (1nM, 5nM, 10nM, 25nM and 50nM) for 1 hour after which 750 ng of Cas9 and 250 ng of sgRNA expression plasmids were transfected with 2 ml of lipofectamine 2000 reagent (Invitrogen) into HeLa cells. After 72 hours from transfection, the cells are detached for genomic DNA extraction. Since the chemical was treated to the cells, the cells were maintained with chemical-containing media.

### Microhomology-dependent deletion events

The alignment information about deletion reads was extracted from alignment results SAM files from k-mer alignment analysis program. Deletion position and the near sequence were calculated using the alignment information. The homology length was calculated by comparing the both sequences in 1bp increments from the start or end position of the deletion sites. If the homology length is 2-16 bp, the reads were categorized as microhomology-dependent.

### Transfection for base editing and prime editing

HEK293T cells (1 x 10^5^ cells per well) were cultivated in a 24-well plate for 24 hours. A mixture of 0.5 ml jetOPTIMUS reagent (Polyplus, 101000006), 500ng plasmid DNA (375ng BE expression plasmid and 125 ng sgRNA expression plasmid) or 543 ng plasmid DNA (365 ng PE expression plasmid, 125 ng pegRNA expression plasmid and 43 ng ngRNA expression plasmid) were added to the cells. After 72 hours, gDNA was isolated.

## Data availability

High-throughput sequencing data have been deposited in the NCBI Sequence Read Archive database (SRA; https://www.ncbi.nlm.nih.gov/sra) under accession number PRJNA1055687.

